# Mechanism of action of methotrexate against Zika virus

**DOI:** 10.1101/481440

**Authors:** Sungjun Beck, Jean A. Bernatchez, Zhe Zhu, Michelli F. Oliveira, Davey M. Smith, Jair L. Siqueira-Neto

**Affiliations:** Skaggs School of Pharmacy and Pharmaceutical Sciences, University of California, San Diego, La Jolla, California, USA 92093; Center for Discovery and Innovation in Parasitic Diseases, University of California, San Diego, La Jolla, California, USA 92093; Sanford Consortium for Regenerative Medicine, La Jolla, California, USA 92093; Department of Medicine, Division of Regenerative Medicine, School of Medicine, University of California, San Diego, La Jolla, California, USA 92093; Department of Medicine, Division of Infectious Diseases, University of California, San Diego, La Jolla, California, USA 92093; Veterans Affairs San Diego Healthcare System, San Diego, California, USA 92093

**Keywords:** Zika virus, antivirals, nucleotide metabolism, methotrexate, dihydrofolate reductase

## Abstract

Zika virus (ZIKV), which is associated with microcephaly in infants and Guillain-Barré syndrome, reemerged as a serious public health threat in Latin America in recent years. Previous high-throughput screening (HTS) campaigns have revealed several potential hit molecules against ZIKV, including methotrexate (MTX), which is clinically used as an anti-cancer chemotherapy and anti-rheumatoid agent. We studied the mechanism of action of MTX against ZIKV in relation to its inhibition of dihydrofolate reductase (DHFR) *in vitro* using Vero and human neural stem cells (hNSCs). As expected, an antiviral effect for MTX against ZIKV was observed, showing up to ten-fold decrease in virus titer during MTX treatment. We also observed that addition of leucovorin (a downstream metabolite of DHFR pathway) rescued the ZIKV replication impaired by MTX treatment in ZIKV-infected cells, explaining the antiviral effect of MTX through inhibition of DHFR. We also found that addition of adenosine to ZIKV-infected cells was able to rescue ZIKV replication inhibited by MTX, suggesting that restriction of *de novo* synthesis adenosine triphosphate (ATP) pools suppresses viral replication. These results confirm that the DFHR pathway can be targeted to inhibit replication of ZIKV, similar to other published results showing this effect in related flaviviruses.

## 1. Introduction

Mosquito-borne flaviviruses, such as Dengue virus (DENV), yellow fever virus (YFV), West Nile virus (WNV), Japanese encephalitis virus (JEV), and Zika virus (ZIKV), are well characterized as human pathogenic viruses transmitted by *Aedes* spp. Mosquitoes [1]. Recently, ZIKV has become a global public health threat that has brought substantial economic burden to affected countries during the recent outbreak in Latin America [2]. Known for its unique neurotropism, ZIKV was associated with a substantial number of ZIKV-induced neurological disorders, mainly microcephaly in infants and Guillain-Barre syndrome, during 2015 and 2016 in Brazil and Colombia [3, 4]. Currently, there are no specific antivirals or vaccines to treat ZIKV infection.

High-throughput screening (HTS) of various compound libraries against ZIKV have been published, and a number of hit molecules against ZIKV were identified [5-8]. One active compound was methotrexate (MTX), which is used to treat a variety of diseases including leukemia, psoriatic arthritis, and rheumatoid arthritis [9-11]. As a chemotherapy agent, the main function of MTX is to antagonize dihydrofolate reductase (DHFR) to decrease *de novo* synthesis of purines and pyrimidines [12, 13], thereby inhibiting the viability of highly replicating cells. Also, MTX has been reported to interfere with a variety of cellular mechanisms, such as oxidative stress or cellular differentiation via methylation [14, 15].

Interestingly, DHFR inhibitors have also been investigated to treat infectious diseases. Due to its formation of a specific dimer with thymidylate synthase (TS) [16], protozoan DHFR has been studied as a drug target to treat parasitic diseases[17]. As antiviral agents, MTX and the antimetabolite floxuridine have been shown to decrease the replication of DENV through inhibition of host DHFR and TS enzymes [18]. Here, we demonstrate the antiviral activity of MTX against ZIKV and probe its mechanism of action using downstream metabolites of the DHFR pathway, leucovorin and adenosine.

## 2. Materials and Methods

### 2.1 Cell and Virus Culture

Vero cells (ATCC CCL-81) were purchased from ATCC and cultured in Dulbecco’s Modified Eagle Medium (DMEM) (Gibco) with high glucose (4.5g/L), 10% fetal calf serum (FCS) (Sigma), and 1% Penicillin-Streptomycin (PS) (Sigma-Aldrich). Human neural stem cells (hNSCs) (Y40050) were purchased from Clontech and cultured in Neurobasal-A medium without phenol red (Thermo Fisher) with the addition of B27 supplement (1:100, Thermo Fisher, #12587010), N2 supplement (1:200, Invitrogen, #17502-048), 20 ng/ml FGF (R&D Systems 4114-TC-01M), 20 ng/ml EGF (R&D Systems 236-EG-01M), GlutaMax (Thermo Fisher, #35050061), and sodium pyruvate. Both Vero and hNSCs were cultured in T75 flasks (Corning). Two strains of ZIKV from Puerto Rico (PRV) (PRVABC59, NR-50240) and Panama (HPAN) (H/PAN/2016/BEI-259634, NR-50210), were cultured on Vero cells in DMEM with high glucose (4.5g/L), 1% fetal bovine serum (FBS) (Sigma), and 1% PS (Sigma-Aldrich). When 100% confluency of Vero cells was reached, the initial medium was removed, and the cells were infected with ZIKV at multiplicity of infection (MOI) of 0.5. At 24h post-infection (PI), the initial medium was removed, and fresh medium was added. At 48h PI, the medium was collected and spun down at 1500 rpm for 10 minutes. The supernatant was collected, 1% DMSO was added, and the sample was stored at −80°C. The titer of the virus stock was measured by plaque assay.

### 2.2 Plaque Assay

Vero cells were seeded at a density of 30,000 cells/well on a 96-wells plate (Corning) and incubated at 37°C and 5% CO_2_ for 24h before infection. Each ZIKV sample was diluted by ten-fold serial dilution, added to Vero cells in at least duplicate, then incubated for 1h. After incubation, media with virus dilutions were removed from the cells, and ZIKV-infected Vero cells were covered with an overlay consisting of DMEM with 0.35% UltraPure agarose (Life Sciences). The plates were further incubated for 72h and fixed with 37% formaldehyde overnight. After fixation, the agarose overlay was aspirated, and fixed cell monolayer was stained with 0.25% crystal violet. For each sample, the titer of ZIKV was reported in PFU/mL.

### 2.3 Immunofluorescence Imaging

Vero cells were seeded at a density of 30,000 cells/well in an opaque black 96-wells plate (Corning) and incubated at 37°C and 5% CO_2_ for 24h before infection. Then, Vero cells were infected with ZIKV at MOI of 0.2. The infected Vero cells were incubated for 48h then fixed with 4% formaldehyde in phosphate buffered saline (PBS). After fixation, the cells were permeabilized with 0.2% TritonX-100 in PBS for 5 min, then blocked with 1% bovine serum albumin (BSA) (Amresco) in PBS for 30 min at room temperature. After blocking, the cells were incubated with diluted primary antibody, anti-flavivirus group antigen primary antibody, clone D1-4G2-4-15 (Millipore #MAB10216)), in PBS with 1% BSA and 0.1% TritonX-100 at 4°C overnight. The cells were washed with PBS three times and treated with diluted secondary antibody, goat anti-mouse Alexa Fluor 594 (ThermoFisher Scientific #A-11032), in PBS with 1% BSA for 1h at room temperature while covered with aluminum foil. The cells were then washed with PBS three times and stained with SYTOX Green (ThermoFisher Scientific #R37168) to visualize nuclei. Immunofluorescence images of cells were acquired using a ZEISS Fluorescence Microscope (Carl Zeiss, Jena, Germany, Axio Vert A1), and images were taken using Zen2 Software. The images were processed with ImageJ (https://imagej.nih.gov/ij/).

### 2.4 Cytotoxicity and Efficacy Study of MTX

The cytotoxic concentration 50 (CC50) of MTX (Sigma #M9929) was determined using two reagents, CellTiter-Glo® (CTG) (Promega #G7570), measuring ATP levels as a readout for cell viability, and CellTiter-Fluor™ (CTF) (Promega #G6080), measuring live-cell protease activity, as per manufacturer’s protocol. To validate the CC50 value from the two cell viability assays, a direct cell number count was performed using trypan-blue staining after MTX treatment. The efficacy of MTX against ZIKV was studied by reduction of ZIKV titer using four different MTX concentrations, 50.0μM, 6.25μM, 0.781μM, and 0.0977μM, in dose-response manner.

### 2.5 Rescue of ZIKV Replication after MTX Treatment by Leucovorin

To test the rescue effect of leucovorin on ZIKV replication with MTX treatment, Vero cells and hNSCs were seeded in a 96-wells plate (Corning) at a cell density of 30,000 and 10,000 cells/well, respectively. The Vero cells and hNSCs were then infected with ZIKV at MOI 0.2 and 0.1, respectively, and simultaneously treated with 5μM MTX, or a combination of 5μM MTX with 50μM folic acid (Sigma #F7876) or leucovorin (Sigma #F7878). The cells were further incubated for 48h at 37°C and 5% CO_2_. After incubation, the supernatant was collected to measure the virus titer. The cells were then treated under the same conditions as described above but without ZIKV infection to assess the effect of leucovorin on countering the cytotoxic effects of MTX on the cells. Then, the cell viability was measured using CTG (Promega) reagent.

### 2.6 Rescue Effect of GAT Medium on ZIKV Replication and Cell Viability from Methotrexate

To test the rescue effect of GAT medium (Glycine, Adenosine, and Thymidine, Sigma #410225, #A4036 and #T1895, respectively) on MTX-mediated suppression of ZIKV replication, Vero cells were seeded in a 96-wells plate at a cell density of 10,000 cells/well. GAT medium was prepared as described previously [19], in MEM (Gibco) with 10% FCS (Sigma) and 1% PS (Sigma-Aldrich) with a final concentration of 0.67mM glycine, 37.5μM adenosine and 41.3μM thymidine. Vero cells were infected with ZIKV at a MOI 0.2 and simultaneously treated with 5μM MTX in GAT medium. The cells were further incubated for 48h at 37°C and 5% CO_2_. After incubation, the supernatant was collected to measure the virus titer. Cells were treated under the same conditions as described above but without ZIKV infection as a cell viability control. Cell viability was measured using CTG reagent (Promega). To investigate the rescue effect of individual components of GAT medium, Vero cells were infected with ZIKV as described above. Then, a combination of 5μM MTX with 0.67mM glycine, 37.5μM adenosine, 41.3μM thymidine, or both adenosine and thymidine were simultaneously added to the infected Vero cells. The treated Vero cells were further incubated for 48h, and supernatant from each sample was collected to measure the virus titer. The same conditions were also applied to Vero cells without ZIKV infection to study which component of GAT medium could protect cells from MTX-induced cytotoxicity, measure by cell viability using both CTG and CTF reagents.

## 3. Results

### 3.1 Inhibition of ZIKV Replication with MTX

To validate the antiviral effect of MTX against ZIKV, immunofluorescence imaging and plaque assay were performed. Through immunofluorescent detection, reduced ZIKV (HPAN MOI 0.2) envelope protein synthesis in Vero cells was observed from 5μM MTX treatment at 48h PI (Figure 1A). We next tested the replication kinetics of ZIKV (HPAN and PRV; both MOI 0.2) in Vero cells by plaque assay at three different time points, 1h, 48h, and 96h PI (Figure 1B, C). Upon 5μM MTX treatment, the replication of two ZIKV isolate strains had greatest reduction (approximately ten-fold decrease in ZIKV titer) at 48h PI, which is consistent with the immunofluorescence images. However, the ZIKV replication resumed at 96 PI, compared to DMSO control, indicating MTX could not continuously suppress ZIKV virion formation after 48h PI. In the case of efficacy of MTX against ZIKV, four different MTX concentrations, 50.0μM, 6.25μM, 0.781μM, and 0.0977μM, were incubated with ZIKV-infected Vero cells (HPAN MOI 0.2) (Figure 1D). At 48h PI the ZIKV titer decreased about ten-fold at 6.25μM MTX treatment.

**Figure 1.**
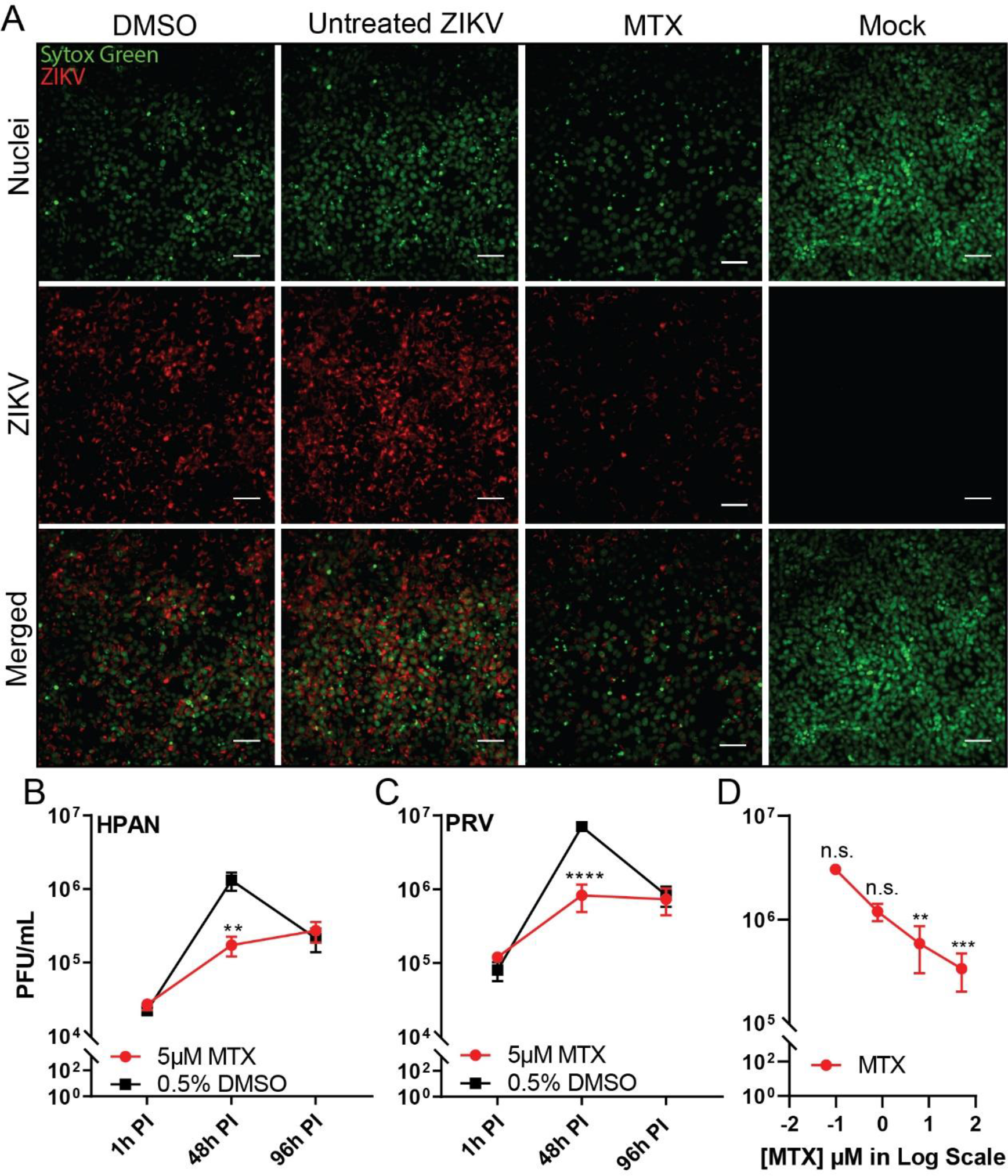
Inhibition of ZIKV replication in Vero Cells after MTX treatment. ZIKV-infected Vero cells were treated with 5μM MTX or 0.5% of DMSO as a negative control. (**A**) Immunofluorescence images (20X) of ZIKV-infected (HPAN MOI 0.2) Vero cells were acquired to analyze the level of ZIKV-Envelope protein after 5μM MTX treatment at 48h PI. SYTOX Green was used to stain nuclei of Vero cells. Scale bars represents 5μm. The virus titer of two ZIKV strains, (**B**) HPAN MOI 0.2 and (**C**) PRV MOI 0.2 were measured after 5μM MTX treatment at three different time points, 1h PI, 48h PI, and 96h PI. (**D**) ZIKV titer after MTX treatment in dose-response manner, 50μM, 6.25μM, 0.781μM, and 0.0977μM, from ZIKV-infected (HPAN MOI 0.2) Vero cells at 48h PI. At least two independent replicates were performed. For (**B**) and (**C**), Two-way ANOVA, followed by Sidak’s multiple comparisons test, were used for statistical analysis. For (**D**), One-way ANOVA, followed by Tukey’s multiple comparisons test, were used for statistical analysis; average viral titer at each MTX concentration was compared to that of untreated ZIKV at 48h PI as a control. Error bars represent standard error of the mean (SEM). ** P ≤ 0.01, *** P ≤ 0.001, **** P ≤ 0.0001, n.s. not significant

### 3.2 Cell Cytotoxicity and Antiviral Efficacy of Methotrexate

To study the cytotoxicity of MTX, we used CTG and CTF reagents for cell viability assays. We tested the compound in different concentrations to calculate CC_50_ of MTX in the two host cells (Table 1). Using CTG reagent, the CC_50_ of MTX in Vero cells and hNSCs were calculated to be 0.104μM and 0.0163μM, respectively. However, using CTF reagent, the CC50 of MTX in Vero cells and hNSCs were calculated to be both >100μM. The two CC50 values showed drastic variance. Accordingly, to validate which CC_50_ value is more appropriate, the number of Vero cells were directly counted under microscopy with trypan-blue staining after 0.5μM of MTX treatment (Figure 2A, B). Although the CC_50_ of MTX using CTG resulted in a value < 0.5μM, the actual number of Vero cells after 0.5μM MTX treatment did not decrease but rather resulted in slowed cellular growth, supporting the CC_50_ (>100μM) value obtained using the CTF reagent.

**Table 1.** CC_50_ values of MTX in Vero cells and hNSCs.

| CC<sub>50</sub>(µM) Values of MTX |  |  |  |
| --- | --- | --- | --- |
|  |  | Vero cells | hNSCs |
| Assay Reagent | CTG | 0.104 | 0.0163 |
|  | CTF | >100 | >100 |

CTG: CellTiter-Glo® measures ATP as a luminescent readout of cell viability biomarker.
CTF: CellTiter-Fluor™ measures live protease activity as a fluorescent readout of cell viability biomarker.
95% CI(µM) of CC<sub>50</sub> value of MTX in Vero cells and hNSCs is 0.0931 - 0.118 and 0.0148 - 0.0181, respectively.

**Figure 2.**
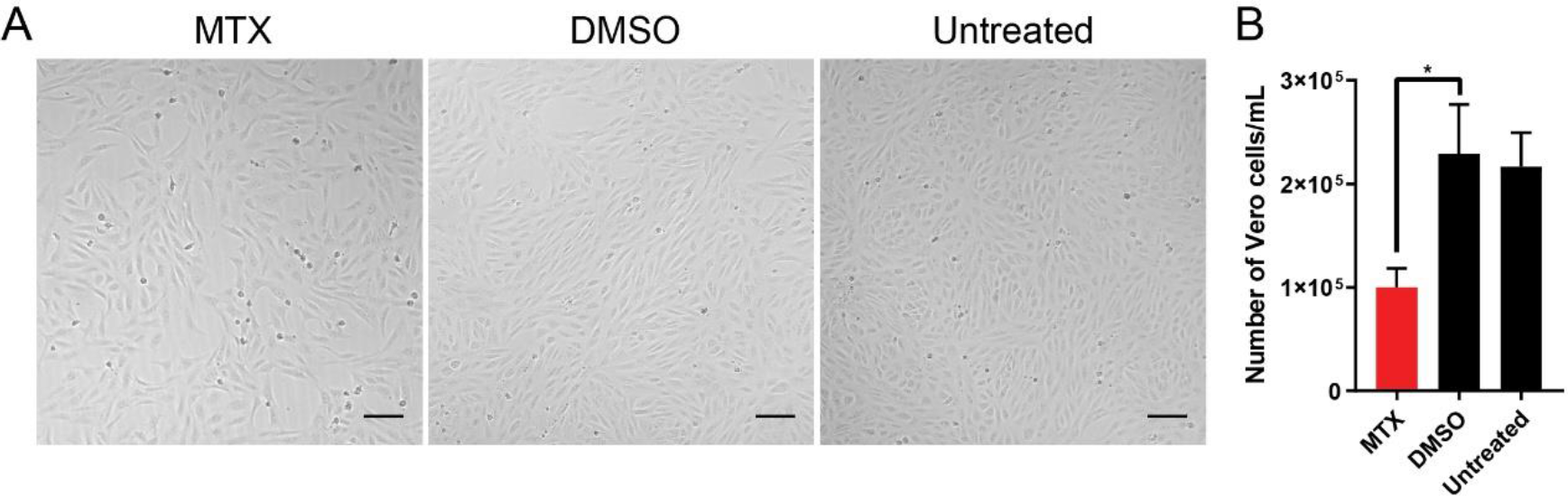
Determination of cytotoxicity of MTX and efficacy of MTX against ZIKV. To determine the cytotoxicity of MTX, the number of Vero cells after MTX treatment was directly counted by trypan-blue staining. Then, the number of Vero cells after MTX treatment was directly counted by trypan-blue staining. Then, the number of live cells were counted under a light microscope with a hemocytometer. (A) 10X bright-field images of Vero cells with 0.5μM MTX treatment. Scale bars represent 5μm. (B) The cell density of Vero cells after 0.5μM MTX treatment after incubation for 48h at 37°C and 5% CO_2_. At least two independent replicates were performed. One-way ANOVA, followed by Tukey’s multiple comparisons test, were used for statistical analysis. The error bars represent the standard error of the mean (SEM). * P ≤ 0.05

### 3.3 Relationship between MTX, Folate, and ZIKV Replication

To understand the mechanism of action for the antiviral effect of MTX, we proceeded to probe the dihydrofolate reductase (DHFR) pathway. DHFR, which reduces dihydrofolate (DHF) to tetrahydrofolate (THF), is a key enzyme for *de novo* synthesis of purines and thymidylate, thereby playing a critical role in cellular growth. As an antifolate, MTX has been reported as a competitive inhibitor of DHFR [20] and shares a similar chemical structure to folate, the natural substrate of DHFR. Through MTX-mediated antagonism of DHFR, folate metabolism is inhibited, which in turn decreases cellular replication. However, addition of leucovorin (folinic acid), a downstream metabolite of the DHFR pathway, can readily rescue the antagonistic effect of MTX in cancer cells [21].

To understand the metabolic relationship between MTX, folate, and leucovorin, Vero cells were infected with two ZIKV isolate strains, HPAN and PRV (both MOI 0.2) and treated with small molecules under three different conditions: MTX, MTX with folic acid, and MTX with leucovorin. At 48h PI, the combination treatment of MTX with folic acid could not rescue the ZIKV replication; however, the combination treatment of MTX with leucovorin rescued ZIKV replication, compared to DMSO control (Figure 3A, B). To study the effect of these three different treatment conditions on Vero host cells, the same experiment was performed with uninfected cells. Using CTG reagent, we measured the cell viability of Vero cells at each condition (Figure 3D). Interestingly, 50μM leucovorin could rescue the cell viability of Vero cells after MTX treatment. Furthermore, the same rescue effect for leucovorin on ZIKV replication and cell viability during MTX treatment was observed in hNSCs (Figure 3C, E). These results support a model where the antiviral effect of MTX against ZIKV occurs through antagonism of DHFR.

**Figure 3.**
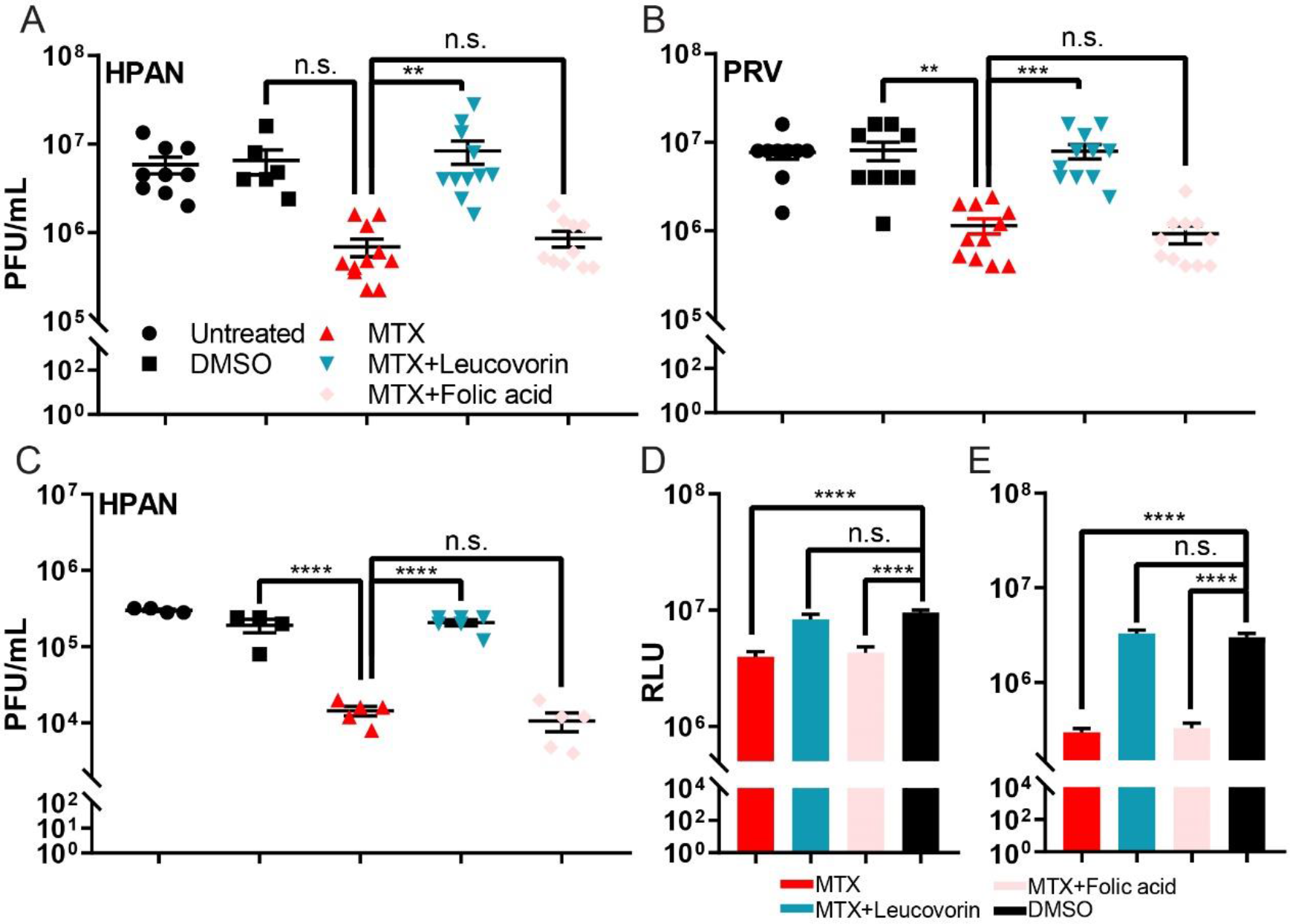
Rescue effect of leucovorin on the cell viability and ZIKV replication during MTX treatment. Two host cell lines, Vero and hNSCs, were used to examine the mechanism of action of MTX against ZIKV replication through the DHFR pathway. Virus titers of the two ZIKV strains, HPAN MOI 0.2 (A) and PRV MOI 0.2 (B) on Vero cells were measured by standard plaque assay. With HPAN infection, the ZIKV titer of the DMSO control was not significantly different from that of 5μM MTX-treated sample in Vero cells (P=0.11) (C). Virus titer of HPAN MOI 0.1 on hNSCs was measured by standard plaque assay. (D). The cytotoxicity of 5μM MTX and co-treatment of MTX with 50μM folic acid or leucovorin on Vero cells was studied by CTG reagent. (E). The cytotoxicity of 5μM MTX and co-treatment of MTX with 50μM folic acid or leucovorin on hNSCs was cells was studied by CTG reagent. RLU relative luminescence unit. At least two independent replicates were performed. One-way ANOVA, followed by Tukey’s multiple comparisons test, was were used for statistical analysis. The error bars represent the standard error of the mean (SEM). ** P ≤ 0.01, *** P ≤ 0.001, **** P ≤ 0.0001, n.s. not significant

### 3.4 Rescue effect of GAT medium on ZIKV Replication and Cell Viability during MTX Treatment

Based on our evidence that the antiviral effect of MTX occurs through antagonism of DHFR, we next assessed which of the metabolites involved in purine and pyrimidine synthesis could affect ZIKV replication. Previous reports have indicated that the cytotoxicity of MTX was reversed by not only leucovorin but also by GAT medium, which contains glycine, adenosine, and thymidine, in human fibroblasts [19]. We therefore studied the rescue effect of GAT medium and each individual component of the medium on ZIKV replication during MTX treatment in Vero cells. GAT medium could rescue both cell viability and ZIKV replication (HPAN MOI 0.2) during MTX treatment (Figure 4A, B). Subsequently, glycine, adenosine, thymidine, and a combination of adenosine and thymidine were individually added with MTX to ZIKV infected Vero cells. Interestingly, only adenosine rescued ZIKV replication during MTX treatment (Figure 4C). To observe if adenosine alone could rescue cell viability during MTX treatment, the cell viability of Vero cells was studied using both CTG and CTF reagents. Whereas glycine and thymidine could not rescue the cell viability, adenosine alone could, in both CTG and CTF assays. (Figure 4D, E).

**Figure 4.**
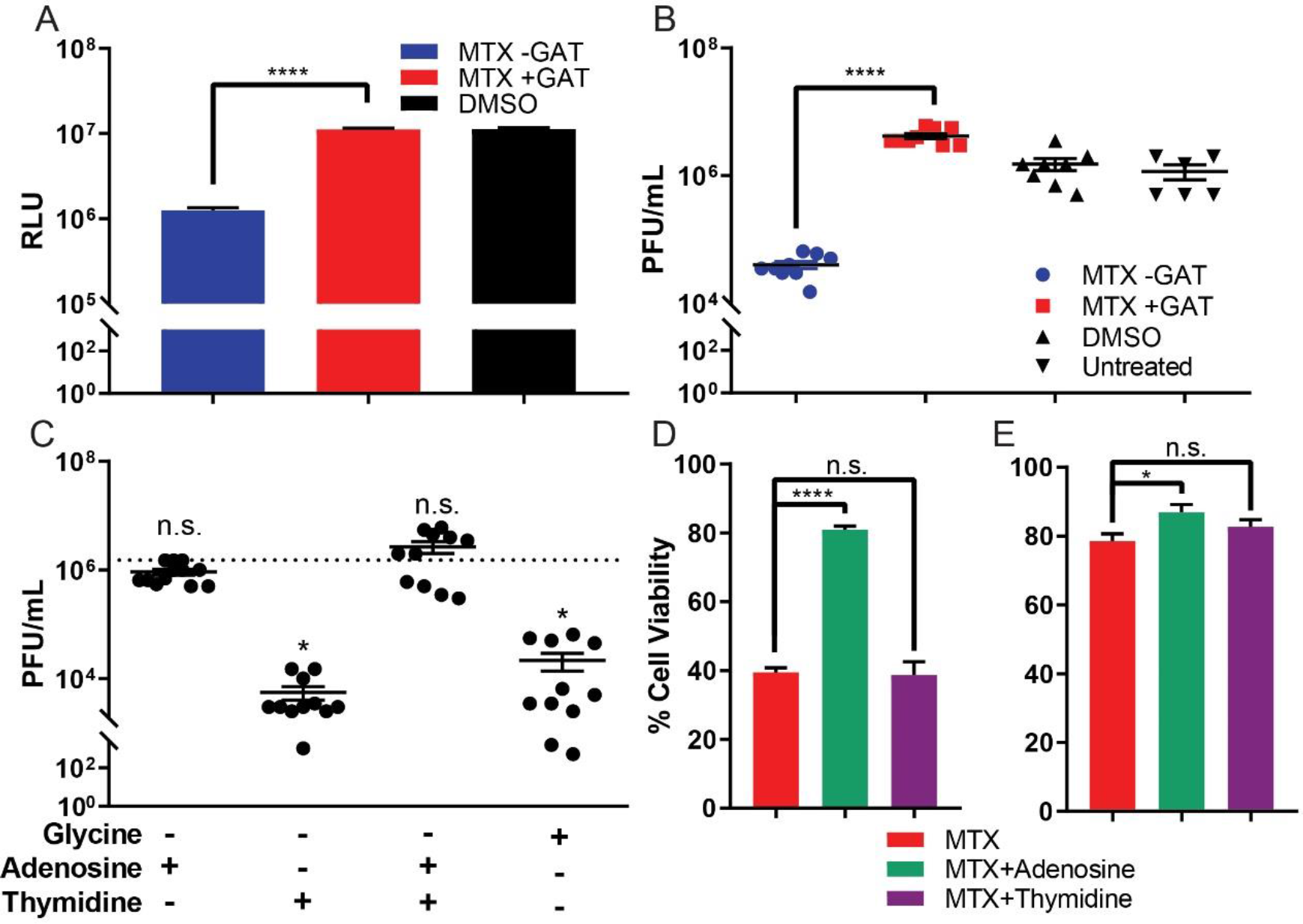
Rescue effect of GAT medium on cell viability and ZIKV replication during MTX treatment. (**A**) Cell viability of Vero cells was studied by CTG reagent after MTX treatment with or without GAT medium. (**B**) GAT medium rescued the ZIKV replication from MTX treatment of the ZIKV-infected (HPAN MOI 0.2) Vero cells. (**C**) Adenosine alone can save ZIKV replication during MTX treatment of the ZIKV-infected (HPAN MOI 0.2) Vero cells. (**D**) Exogenous adenosine rescued cellular ATP levels, but thymidine and glycine could not rescue ATP levels during MTX treatment in Vero cells as measured by CTG assay. (**E**) Adenosine also rescued live-protease activity measured by CTF reagent during MTX treatment in Vero cells. RLU relative luminescence unit. At least two independent replicates were performed. The error bars represent the standard error of the mean (SEM). One-way ANOVA, followed by Tukey’s multiple comparisons test, was used for statistical analysis. * P ≤ 0.05, **** P ≤ 0.0001, n.s. not significant

## 4. Discussion

In this study, we validated MTX as an antiviral targeting ZIKV replication. MTX was previously discovered as a potent hit molecule against ZIKV in human brain microvascular endothelial cells, with inhibitory concentration 50 (IC_50_) of 0.28μM and host cytotoxicity (CC50) of >10μM [5]. Such neurotropism may explain the possible neurological consequences of ZIKV infection, and that MTX influences the folate pathway may explain that one of these consequences is a neural tube defect, i.e. microencephaly in infants. MTX is approved by the FDA to treat diseases such as cancer and rheumatoid arthritis. Accordingly, the mechanism of action of MTX against such diseases have been well studied. Primarily, MTX is known to antagonize DHFR as a competitive inhibitor [20] and antagonized DHFR causes inhibition of *de novo* synthesis of purines and pyrimidines, which are essential for cell replication [22]. Hence, MTX shows great cytotoxicity to highly replicating cancer cell lines. Interestingly, with 5μM MTX treatment, reduced ZIKV replication was observed from infected Vero cells and hNSCs. Although MTX reduced ZIKV titer about ten-fold at 48h PI, MTX could not continuously suppress the ZIKV replication after 48h PI. Observing the antiviral effect of MTX against ZIKV, the toxicity of MTX in Vero cells and hNSCs were measured. Understanding that MTX can inhibit purine and pyrimidine metabolism, cells treated with MTX showed significantly decreased viability in terms of ATP level. Accordingly, CC_50_ measurement with CTG reagent also resulted in high cytotoxicity in the two host cells; however, such toxicity could be misleading because the drug mechanism directly interferes with ATP levels, i.e. the readout measurement of the assay. Accordingly, Vero cells with 0.5μM MTX treatment, exceeding the CC50 value obtained from CTG reagent, did not exhibit phenotypic cytotoxicity but rather showed slower cellular replication compared to untreated controls.

MTX decreased ZIKV titer in Vero cells and hNSCs by antagonizing DHFR. The inhibited DHFR activity by MTX was readily reversed by simultaneous co-treatment with leucovorin, thereby rescuing ZIKV replication from MTX as well. Further metabolite analysis with GAT medium allowed us to understand adenosine alone could rescue ZIKV replication by increasing ATP level from MTX treatment. Although MTX is known to inhibit TS [23], thymidine alone could not rescue the ZIKV replication. Considering the genome of ZIKV, a single stranded positive sense RNA (ssRNA+), thymidine triphosphate (TTP) would not be incorporated into its genome during the replication. In fact, Fischer *et al.* reported inhibition of TS can reduce flavivirus replication but also found DENV replication was not rescued when excess amount of thymidine was co-treated with MTX, suggesting an alternative mechanism such as activation of P53 for general antiviral activity due to low levels of thymidine [18].

Together, our results identified that DHFR may be a promising target for antiviral research against ZIKV and other flaviviruses, or Chikungunya virus that shares similar features to those of ZIKV [24], and this should be further investigated. However, repurposing MTX as an antiviral against ZIKV infection would not be ideal option because the most vulnerable ZIKV infected patients are pregnant women and MTX has been used as a clinical abortion agent [25]. Furthermore, although excess amount of thymidine could not rescue the cell viability from MTX *in vitro* (Figure 4D, E), such combination should be tested *in vivo*. Considering the number of reports on the rescue effect of thymidine from MTX [26-28], perhaps, MTX-thymidine could be an optional medical regime to treat ZIKV infection: while MTX inhibits ZIKV replication, thymidine would rescue the host from the cytotoxicity of MTX. In conclusion, this study clarified the antiviral mechanism of MTX, and suggests the DHFR pathway may be targeted for antiviral purpose.

## Author Contributions

S.B., J.A.B., and J.L.S.N. conceived and designed the experiments; S.B. performed experiments; S.B., J.A.B., Z.Z., M. F. O., D. M. S., and J.L.S.N. analyzed the data; S.B. wrote the original draft of manuscript; J.A.B., M. F. O., D. M. S., and J.L.S.N. reviewed and edited the manuscript.

## Funding

This research was supported by Clinical and Translational (CTRI) pilot grant UL1TR001442 (to J.L.S.N.) and by NIH grants AI100665 and AI036214 (to D.M.S.).

## Acknowledgments

The authors would like to give special thanks to Dr. Alex E. Clark for his great help in preparing ZIKV sample stocks and for sharing reagents for the project.

## Conflicts of Interest

The authors declare no conflict of interest.

---

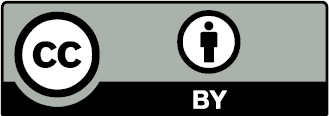

© 2018 by the authors. Submitted for possible open access publication under the terms 377 and conditions of the Creative Commons Attribution (CC BY) license 378 (http://creativecommons.org/licenses/by/4.0/).

